# Structural basis of ion – substrate coupling in the Na^+^-dependent dicarboxylate transporter VcINDY

**DOI:** 10.1101/2022.01.11.475879

**Authors:** David B. Sauer, Jennifer J. Marden, Joseph C. Sudar, Jinmei Song, Christopher Mulligan, Da-Neng Wang

**Author notes:** Correspondence to: C.M., D.N.W. Centre for Medicines Discovery, Nuffield Department of Medicine, University of Oxford, Oxford, UK.

## Abstract

The Na^+^-dependent dicarboxylate transporter from *Vibrio cholerae* (VcINDY) is a prototype for the divalent anion sodium symporter (DASS) family. While the utilization of an electrochemical Na^+^ gradient to power substrate transport is well established for VcINDY, the structural basis of this coupling between sodium and substrate binding is not currently understood. Here, using a combination of cryo-EM structure determination, succinate binding and site-directed cysteine alkylation assays, we demonstrate that the VcINDY protein couples sodium- and substrate-binding via a previously unseen induced-fit mechanism. In the absence of sodium, substrate binding is abolished, with the succinate binding regions exhibiting increased flexibility, including HP_in_b, TM10b and the substrate clamshell motifs. Upon sodium binding, these regions become structurally ordered and create a proper binding site for the substrate. Taken together, these results provide strong evidence that VcINDY’s induced-fit mechanism is a result of the sodium-dependent formation of the substrate binding site.

## Introduction

VcINDY is a Na^+^-dependent dicarboxylate transporter that imports TCA cycle intermediates across the inner membrane of *Vibrio cholerae* ^1, 2^. The detailed structural and mechanistic understanding of VcINDY ^1–4^ has made the protein a prototype of the divalent anion sodium symporter (DASS) family (Supplementary Fig. 1a&b)^5^. Within the human genome, the SLC13 genes encode for DASS members including the Na^+^-dependent, citrate transporter (NaCT) and dicarboxylate transporters 1 and 3 (NaDC1 and NaDC3) ^6^. Besides functioning as TCA cycle intermediates, DASS-imported substrates are central to a number of cellular processes. In bacteria C4-carboxylates can serve as the sole carbon source for growth ^7^, while imported citrate and tartrate are electron acceptors during fumarate respiration ^8^. Citrate is also a precursor for both fatty acid biosynthesis and histone acetylation in mammals ^9, 10^. Dicarboxylates such as succinate and *α*-ketoglutarate act as signaling molecules that regulate the fate of naive embryonic stem cells and certain types of cancer cells ^11, 12^. As a result of these roles in regulating cellular di- and tricarboxylate levels, mutations in DASS transporters have dramatic physiological consequences. Deletion of bacterial DASS transporters can abolish growth on particular dicarboxylates ^7, 8^. Mutations in the human NaCT transporter cause SLC13A5-Epilepsy in newborns ^13^, whereas variants in the dicarboxylate transporter NaDC3 lead to acute reversible leukoencephalopathy ^14^. In mice, knocking out NaCT results in protection from obesity and insulin resistance ^15^. Such roles of SLC13 proteins in cell metabolism have made them attractive targets for the treatment against obesity, diabetes, cancer and epilepsy ^16–18^. Therefore, mechanistic characterization of the prototype transporter VcINDY will help us to better understand the transport mechanism of the entire DASS family, including the human di- and tricarboxylate transporters.

The VcINDY protein is a homodimer consisting of a scaffold domain and an elevator domain (Supplementary Fig. 1b-e) ^1^. The conservation of this architecture throughout the DASS/SLC13 family has been confirmed by X-ray crystallography and cryo-electron microscopy (cryo-EM) structures of VcINDY, LaINDY, a dicarboxylate exchanger from *Lactobacillus acidophilus,* and the human citrate transporter NaCT ^1, 4, 19, 20^. Comparison of VcINDY in its inward-facing (Ci) conformation with the outward-facing (Co) structure of LaINDY, along with MD simulations, reveals that an elevator-type movement of the transport domain, through an ∼12 Å translation along with an ∼35° rotation, facilitates translocation of substrate from one side of the membrane to the other ^19^. In fact, the structural and mechanistic conservation may extend beyond DASS to the broader Ion Transport Superfamily (ITS) ^5, 21^.

Substrate transport of VcINDY is driven by the inwardly-directed Na^+^ gradient, with dicarboxylate import coupled to the co-transport of three sodium ions (Supplementary Figs. 1a&b) ^1, 2, 22^. The binding sites for the substrate and two central Na^+^s have been identified in the structures of VcINDY in its Na^+^- and substrate-bound inward facing (Ci-Na^+^-S) state (Supplementary Figs. 1e) ^1, 4^. The Na1 site on the N-terminal half of the transport domain is defined by a clamshell formed by loop L5ab and the tip of hairpin HP_in_. A second clamshell encloses Na2, related to Na1 by inverted-repeat pseudo-symmetry in the sequence and structure, and formed by L10ab and the tip of hairpin HP_out_ (Supplementary Fig. 1c). In addition to binding the Na^+^s, both hairpin tips also form parts of the substrate binding site, located between the Na^+^ sites. Each hairpin tip consists of a conserved Ser-Asn-Thr (SNT) motif, and the two SNT motifs form part of the substrate binding site, making direct contact with carboxylate groups of the substrate. Whereas these two SNT signature motifs are responsible for recognizing carboxylate, additional residues in neighboring loops have been proposed to distinguish between different kinds of substrates ^4^. Notably, VcINDY’s structure with sodium in the absence of a substrate (the C_i_-Na^+^ state), determined in 100 mM Na^+^, is very similar to the substrate-bound state C_i_-Na^+^-S ^19^.

**Fig. 1.**
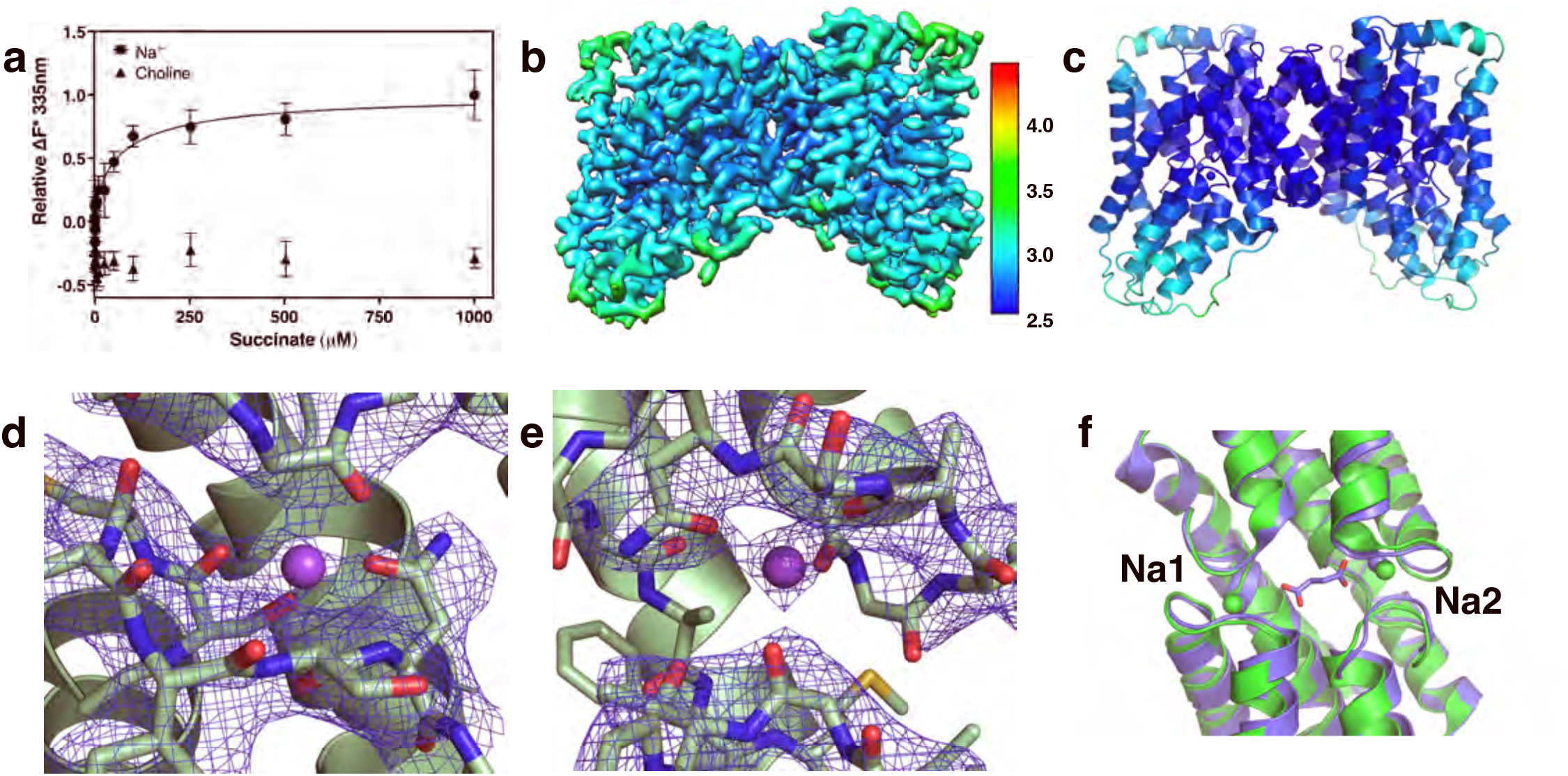
Cryo-EM structure of VcINDY in the C_i_-Na^+^ state determined in 300 mM Na^+^. **a**. Measurements of succinate binding to detergent-purified VcINDY in the presence of 100 mM NaCl, using intrinsic tryptophan fluorescence quenching (*N* = 4). The *K*_d_ was determined to be 92.17± 47.38 μM. When NaCl was replaced with Choline chloride, no binding of succinate to VcINDY could be measured (*N* = 4). **b**. Cryo-EM map of VcINDY determined in the presence of 300 mM NaCl. The map is colored by local resolution (Å). The overall map resolution is 2.83 Å. **c**. Structure of VcINDY in the C_i_-Na^+^ state. The structure is colored by the B-factor. **d**. Na1 site structure and Coulomb map. **e**. Na2 site structure and Coulomb map. **f**. Overlay of VcINDY structures around the substrate and sodium binding sites in the C_i_-Na^+^ state (green) and C_i_-Na^+^-S state (PDB ID: 5UL7, blue). There is very little structural change observed between the two states.

While the Na^+^- and substrate-binding sites in VcINDY have been well-characterized ^1, 4, 23^, the coupling mechanism between the electrochemical gradient and substrate transport ^24^ is less well understood. There is strong evidence that charge compensation by sodium ions is essential to lowering the energy barrier for transporting the di- and trivalent anionic substrates across the membrane ^19^. However, such charge compensation alone does not necessarily result in substrate binding as Li^+^ is able to bind to VcINDY similarly to Na^+^, but results in a lower affinity substrate binding site and considerably reduced transport rates ^2, 23^. More importantly, charge neutralization cannot explain the sequential binding observed for VcINDY. As is the case for other DASS proteins ^25–28^, all available experimental evidence from both whole cells and reconstituted systems supports the notion that in VcINDY sodium ions and substrate bind in a sequential manner, with Na^+^s binding first, followed by dicarboxylate ^2, 3, 23, 29^. As a secondary-active transporter can transport substrate in either direction, it follows that the release of the substrate and Na^+^s is also ordered, with the substrates being released first.

Structures of VcINDY in the Na^+^- and substrate-bound state C_i_-Na^+^-S, in which the Na^+^ sites share residues with substrate sites in their center, allowed us to propose that substrate binding in VcINDY follows an induced-fit mechanism ^1^. In this mechanism, the binding of sodium ions helps to create a proper binding site for substrate (Supplementary Fig. 1a&b). Conversely, in the absence of bound sodium ions the substrate-binding site will change significantly, such that the substrate cannot bind. Not only can such a mechanism be part of Na^+^ — substrate coupling, it may also explain the sequential binding observed for VcINDY.

This induced-fit mechanism of substrate binding enables us to make two explicit, experimentally testable predictions. First, the affinity of the transporter to a substrate must be much higher in the presence of Na^+^ than in its absence. Second, substantial structural changes will occur at the Na^+^ sites in the absence of sodium, affecting substrate binding. In this work, we aimed to test these two predictions using a combination of structure determination by single particle cryo-EM, substrate binding affinity measurements by intrinsic tryptophan fluorescence quenching, and position accessibility quantification via a newly-developed site-directed cysteine alkylation assay^29^. These experimental results allow us to directly test the induced-fit binding model of VcINDY.

## Results

### Succinate binding depends on the presence of Na^+^

To test the first of the predictions generated from our induced-fit hypothesis, we measured VcINDY’s binding affinity for the model substrate, succinate, in the presence and absence of Na^+^ (Supplementary Fig. 2). To assess substrate binding we used intrinsic tryptophan fluorescence quenching, which has been successfully applied to measure substrate binding for various membrane transporters ^20, 30–35^. In the presence of 100 mM Na^+^, detergent-purified VcINDY was found to bind succinate with a *K*_d_ of 92.17 ± 47.38 μM (Fig. 1a, Supplementary Fig. 2b). For comparison, the human NaCT in the same protein family binds its substrate citrate at a *K*_d_ of 148 ± 28 μM ^20^.

To measure the binding affinity of succinate to VcINDY with empty Na^+^ binding sites, we searched for a cation to replace Na^+^ in the purification buffer. This ion should not occupy the Na1 or Na2 sites while still allowing the transporter protein to remain stable in solution. K^+^ is unable to power transport by VcINDY, but was found to be unsuitable as the protein precipitated when purified in the presence of 100 mM KCl. We next tested the organic cation choline (C_5_H_14_NO^+^, Ch^+^ in abbreviation). We reasoned that Ch^+^ would be more stabilizing than K^+^ based its position in the Hofmeister series ^36^, and that its size would preclude it from occupying Na^+^ binding sites ^37^. Indeed, VcINDY purified in 100 mM NaCl remained soluble at 0.5 – 1.0 mg/mL after diluting the sample 11,000-fold in 100 mM ChCl. VcINDY was therefore purified in the presence of 100 mM Ch^+^ as the only monovalent cation. The protein eluted as a sharp, symmetrical peak on a size-exclusion chromatography column (Supplementary Fig. 2a), confirming its stability and structural homogeneity.

Notably, intrinsic tryptophan fluorescence quenching with VcINDY purified and assayed in the presence of 100 mM Ch^+^ revealed no succinate binding (Fig. 1a, Supplementary Fig. 2c). Thus, the binding measurements of VcINDY in the presence and absence of Na^+^ are consistent with an induced-fit model where bound sodium ions are necessary to create a proper binding site for succinate. Encouraged by these findings and our ability to produce stable, structurally homogeneous and Na^+^-free VcINDY, we next sought to uncover the structural basis of this Na^+^ — substrate coupling by determining the transporter’s structures using cryo-EM in different states.

### Structure of VcINDY in 300 mM Na^+^

Generally speaking, the transport mechanism of a secondary-active transporter is reversible, in which the direction of substrate translocation depends on direction and magnitude of the driving force (Supplementary Fig. 1a&b). Consequently, substrate binding is equivalent to substrate release. Therefore, to provide structural insights into VcINDY’s binding process, we aimed to characterize the substrate release process in the inward-facing (C_i_) conformations by capturing the structures of VcINDY in the following states: its Na^+^-and substrate-bound state (C_i_-Na^+^-S), its Na^+^-bound state (C_i_-Na^+^) and its Na^+^- and substrate-free state (C_i_-apo).

The C_i_-Na^+^-S structure of VcINDY has previously been solved using X-ray crystallography ^1, 4^. Additionally, we had characterized the C_i_-Na^+^ state using a cryo-EM structure of VcINDY purified in 100 mM Na^+^ without substrate ^19^. The similarity of these structures agrees with our induced-fit model of Na^+^ – substrate coupling, which does not require the structure of the sodium-bound C_i_-Na^+^ state to be significantly different from that of the C_i_-Na^+^-S state with both cations and substrate bound (Supplementary Figs. 1a&b). However, as the apparent *K*_m_ for Na^+^ for VcINDY was measured to be 41.7 mM ^2^, our earlier VcINDY sample in 100 mM Na^+^ likely represents a mixture of the C_i_-Na^+^ and C_i_-apo states. It is unclear whether the subsequent cryo-EM image processing of the particles was able to exclude all particles of the Na^+^-free C_i_-*apo* state. To more clearly and definitely resolve the C_i_-Na^+^ state structure, in the current work we purified and determined a structure of VcINDY in 300 mM Na^+^. This ion concentration was optimized to increase the Na^+^ occupancy while, at the same time, to ensure a low enough noise level in the cryo-EM images to determine a C_i_-Na^+^ state structure of this small membrane protein (total dimer mass: 126 kDa) at 2.83 Å resolution (Fig. 1b,c, Supplementary Figs. 3 & 4a-c, Table 1).

We previously determined two cryo-EM structures of VcINDY in the presence of 100 mM NaCl ^19^. The new structure in 300 mM Na^+^ (Fig. 1c, Supplementary Fig. 4b) is identical to the previously determined one bound to a Fab and embedded in lipid nanodisc in 100 mM Na^+^ (PDB ID: 6WW5) ^19^ (r.m.s.d. of 0.460 Å for all the non-hydrogen atoms), except for the position of the last three residues at the C-terminus, which interact with the Fab molecule used for structure determination (Supplementary Fig. 4d). Furthermore, though the new map clarified the loop connecting HP_out_ b and TM10b, the new model in 300 mM NaCl is effectively identical to the other previous C_i_-Na^+^ structure in 100 mM NaCl, determined in amphipol and without Fab (PDB ID: 6WU3) ^19^, with an r.m.s.d of 0.358 Å after excluding Val392 – Pro400 (Supplementary Fig. 4d).

As expected from the higher Na^+^ occupancy in the 300 mM sample, better defined densities appeared within both the Na1 and Na2 clamshells (Fig. 1d&e), which were absent in the previous 100 mM Na^+^ maps ^19^. In addition to coordination by side chains and backbone carbonyl oxygens, the sodium ion at the Na1 site is stabilized by the helix dipole moments from HP_in_ b and TM5b (Fig. 1f; Supplementary Fig. 1e), as previously observed in other membrane proteins ^38, 39^. Similarly, the Na^+^ ion in the Na2 site is stabilized by HP_out_ b and TM10b.

Finally, this new, higher resolution map confirmed our earlier observations that succinate release caused only limited changes at the substrate binding site without relaxing the two Na^+^ clamshells ^19^. Both the overall structure and the sodium binding sites in the C_i_-Na^+^ state are similar to those in the sodium- and substrate-bound C_i_-Na^+^-S state (Fig. 1f, Supplementary Fig. 4e).

### *Apo* structure of VcINDY in Choline^+^

With the structures of sodium- and succinate-bound ^1, 4^ and Na^+^-only bound (Fig. 1) states in hand, the missing piece of the puzzle to validate the Na^+^ induced-fit mechanism was the C_i_-*apo* state structure of the transporter protein. As a Ch^+^ ion is too large to fit into a Na^+^ binding site ^35, 37^, and VcINDY was stable and monodisperse in the presence of 100 mM ChCl (Supplementary Fig. 2a), such a preparation allowed us to obtain cryo-EM maps of the C_i_-*apo* state (Fig. 2, Supplementary Figs. 5 & 6, Table 1).

**Fig. 2.**
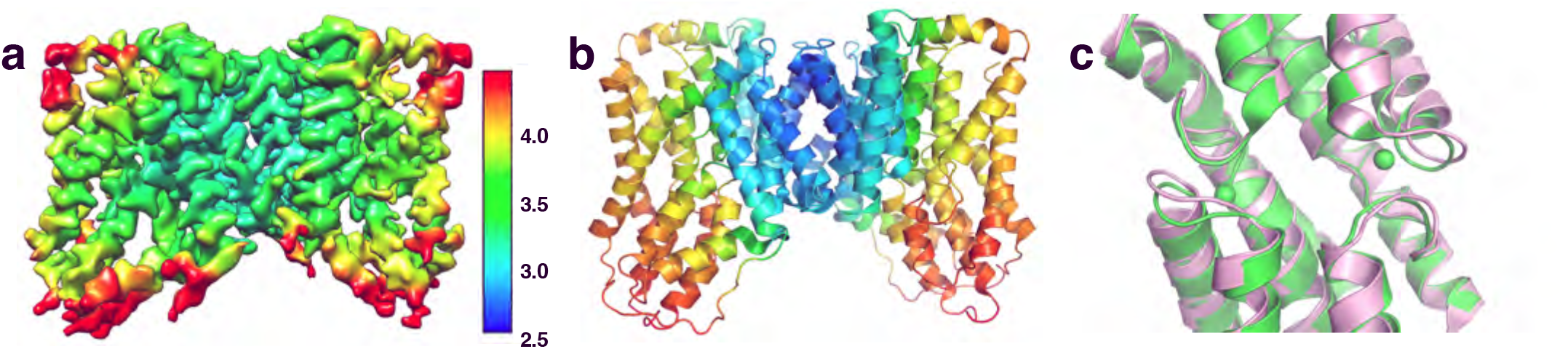
Cryo-EM structure of VcINDY in the C_i_-*apo* state determined in Choline^+^. **a**. Cryo-EM map of VcINDY preserved in amphipol determined in the presence of 100 mM Choline-Chloride. The map is colored by local resolution (Å) on the same scale as Figure 1b. The overall map resolution is 3.23 Å. **b**. Structure of VcINDY in the C_i_-*apo* state. The structure is colored by the B-factor on the same scale as Figure 1b. **c**. Overlay of VcINDY structures around substrate and sodium binding sites in the sodium-bound C_i_-Na^+^ state (green) and the C_i_-*apo* state (pink). The structures of the two Na1 and Na2 clamshells has changed in the absence of sodium ions.

Unlike the VcINDY map in 300 mM Na^+^ for which 3D classification converged to a single map, the VcINDY-choline dataset yielded four classes at a resolution range of 3.6 — 4.4 Å resolution. The 3D class with the highest resolution was further refined to 3.23 Å resolution (Fig. 2a&b, Supplementary Fig. 6c) Whereas the overall fold of the protein in Ch^+^ remains the same (Supplementary Fig. 6d&f), Na^+^ densities within the Na1 and Na2 clamshells are totally absent. Additional local changes are observed for the protein parts near the Na1 and Na2 sites, with a loss of density in each Ci-*apo* map at the HP_in_b and TM10b helices (Fig. 3b), indicating increased local structural flexibility.

**Fig. 3.**
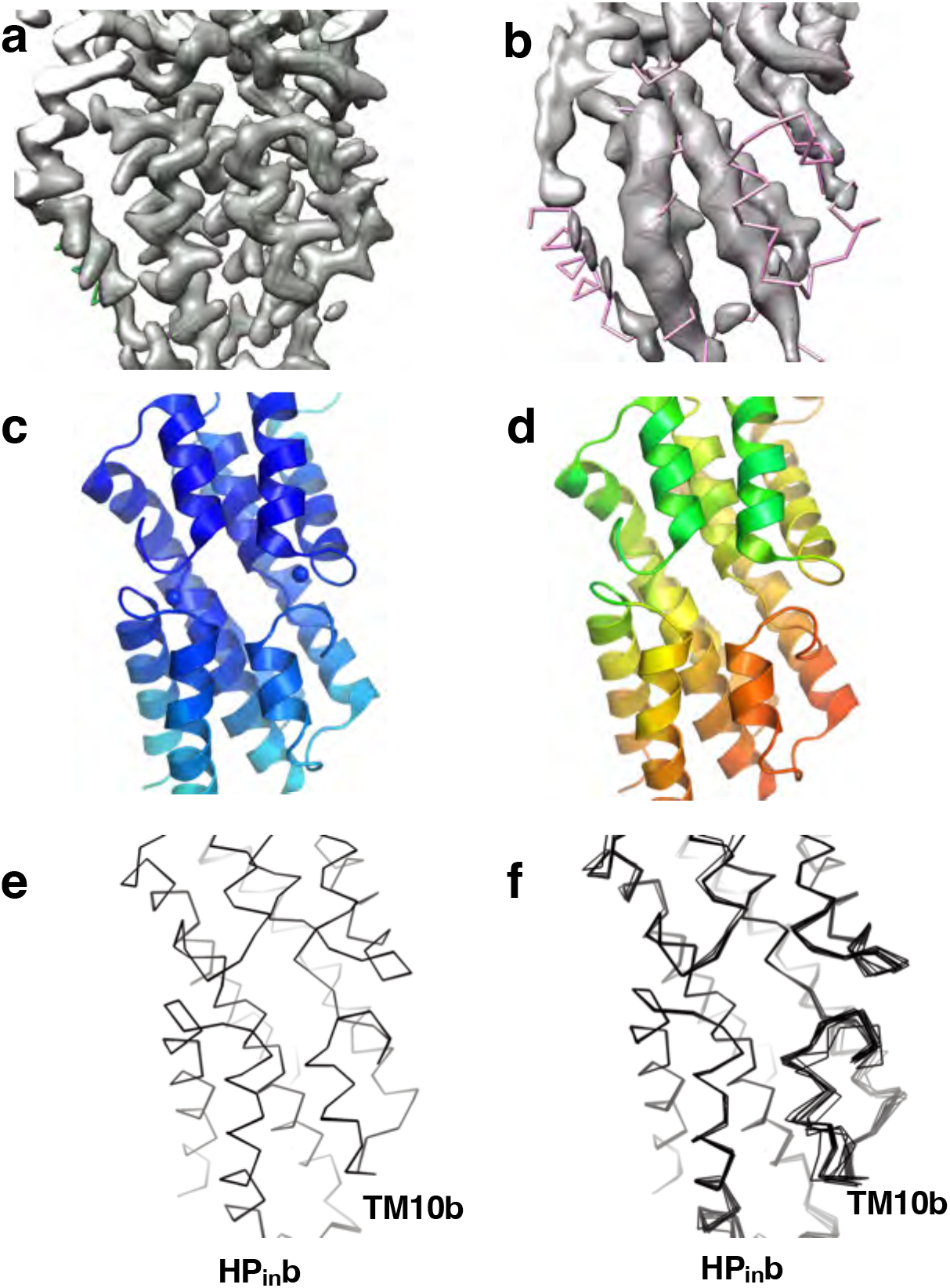
VcINDY flexibility changes near the Na1 and Na2 sites between the C_i_-Na^+^ state and C_i_-*apo* states. **a**. Cryo-EM density map in 300 mM NaCl. **b**. Cryo-EM density map in 100 mM Choline-Chloride. In **a** and **b**, the respective protein models’ backbones are fitted into the densities. **c**. Structure of VcINDY in its C_i_-Na^+^ state. **d**. Structure of VcINDY in its C_i_-*apo* state. In **c** and **d**, the structures are colored by the B-factors on the same scale. **e**. NMR-style analysis of the VcINDY structure in Na^+^. **f**. NMR-style analysis of the VcINDY structure in Choline^+^. The resolution for refinement of both structures in **e** and **f** was truncated to 3.23 Å. In the absence of sodium, the helices on the cytosolic side of Na1 and Na2, particularly HP_in_b and TM10b, show markedly increase flexibility.

### Flexibility of *Apo* VcINDY near the Na1 and Na2 sites

The 3.28 Å VcINDY C_i_-*apo* state structure exhibited significant changes near the Na1 and N2 sites (Figs. 2c, 3a&b), though the overall structure and the average atomic positions are similar to those in the 300 mM Na^+^ (r.m.s.d of 0.672 Å) (Supplementary Fig. 6f). Notably, HP_in_b near the Na1 site and TM10b near the Na2 site showed marked decreased density in the cryo-EM map, corresponding to increased flexibility of this region (Fig. 3a&b). Correspondingly, the model exhibited significantly higher B-factors in the same region compared to the rest of the model (Fig. 3c&d).

To further analyze this flexibility, we used simulated annealing ^40, 41^ in a model refinement protocol analogous to protein structure determination by NMR spectroscopy^42^. We reasoned that in multiple, separate refinements with simulated annealing the rigid parts of the VcINDY would converge to the same coordinates, while mobile portions of the protein would arrive at distinct atomic positions in each run. We term this as NMR-style analysis, though in cryo-EM the constraints being Coulomb potential maps rather than distance constraints.

Most parts of the VcINDY structure exhibit no variation in both the Ci-Na and Ci-apo states, including HP_in_b and TM10b in the 300 mM Na^+^ condition (Fig. 3e, Supplementary Fig. 7a). In contrast, the NMR-style analysis clearly illustrated the structural heterogeneity of the HP_in_b and TM10b regions of the Ci-apo state (Fig. 3f, Supplementary Fig. 7b). The mean r.m.s.d. of the backbone atoms for the Ci-*apo* protomers is 0.150 Å, as opposed to 0.063 Å among C_i_-Na^+^ protomers refined using the same protocol. Such helix flexibility results from the absence of Na^+^ interactions with residues in the clamshells and with the dipoles of HP_in_b and TM10b ^43, 44^.

### Site-directed alkylation supports structural changes to Na1 and Na2 sites

To confirm the local conformational changes and helix flexibility observed in our VcINDY structures, we implemented a site-directed cysteine alkylation strategy that can directly assess the solvent accessibility of specific positions in a protein. In this approach, single cysteines are introduced into a Cys-less version of VcINDY, which is capable of robust Na^+^-driven transport ^2, 3^. Following purification, the cysteine mutants of VcINDY are incubated with the thiol-reactive methoxypolyethylene glycol maleimide 5K (mPEG5K). This mass tag reacts with solvent-accessible cysteines and increases the protein mass by ∼5 kDa, which is separable from unmodified protein on an SDS-PAGE gel. As mPEG5K will react faster with cysteines that are more accessible, monitoring PEGylation over time provides us with the ability to follow changes in accessibility of particular parts of the protein under different conditions ^29^.

To test our induced-fit model, we designed a panel of single-cysteine mutants of VcINDY that would report on the Na^+^-dependent accessibility changes at the Na1 and Na2 sites predicted from structures (Fig. 4a, Supplementary Fig. 7d). We selected residues that, if our mechanistic model is accurate, will exhibit a higher rate of PEGylation in the absence of Na^+^ compared to its presence due to the increased mobility of HP_in_ and TM10b. To create our panel proximal to the Na1 site, we purified five cysteine mutants whose reactive thiol groups are buried in the C_i_-Na^+^ state behind HP_in_ (L138C on HP_in_a, A155C and V162C on HP_in_b and A189C on TM5a). However, similar cysteine substitutions near Na2 (Val427, Ile433, Gly442 and Met438) resulted in diminished expression levels, likely indicating the importance of these residues to the stability of the protein. Fortunately, cysteine mutant of Val441, a residue located on TM11 and behind TM10b (Fig. 4a, Supplementary Fig. 7d), expressed well and allowed for purification. Typically, such single cysteine mutants are capable of Na^+^-driven succinate transport in proteoliposomes ^29^.

**Fig. 4.**
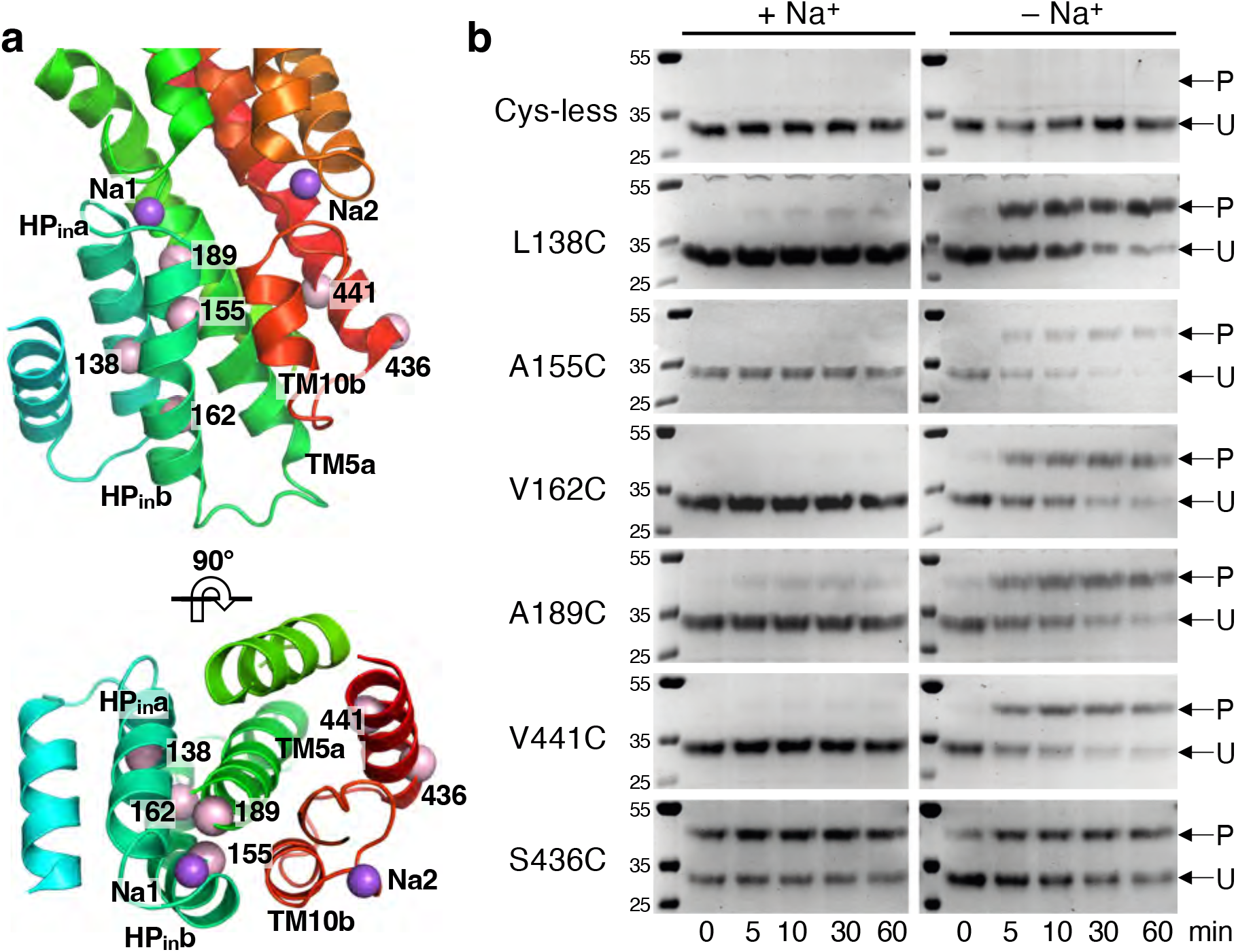
Cysteine alkylation with mPEG5K of VcINDY near the Na1 and Na2 sites in the presence and absence of Na^+^. **a**. Location of cysteine mutations. Our structures suggested that HP_in_b and TM10b become flexible in the absence of sodium, increasing the solvent accessibility of Leu138, Ala155, Val162 and Ala189 near the Na1 site, and Val441 near the Na2 site. Position Ser436, for which no accessibility change was observed between our structures, is used as a control. On a Cys-less background, residues at these positions were individually mutated to a cysteine for mPEG5K labeling. For clarity, only amino acid numbers are labeled and the types are omitted. **b**. Coomassie Brilliant Blue-stained non-reducing polyacrylamide gels showing the site-directed PEGylation of each cysteine mutant over time in the presence and absence of Na^+^. P: PEGylated protein; U: un-PEGylated protein. Each reaction was performed on two separate occasions with the same result.

We monitored the PEGylation of each mutant in the presence and absence of Na^+^. Under these reaction conditions there is no PEGylation of the Cys-less variant demonstrating no background labelling that could hinder analysis (Fig. 4b, top row). In the presence of 300 mM Na^+^ we observed complete inhibition of PEGylation at every position (Leu138, Ala155, Val162, Ala189 and Val411) over the time course of 60 min (Fig. 4b, left panels), in agreement with our model that these residues are buried in the Na^+^-bound state. However, in the absence of Na^+^ (but with 300 mM Ch^+^), every mutant showed escalated levels of PEGylation over time (Fig. 4b, right panels), indicating the increased flexibility of HP_in_b and TM10b.

To ensure that the change in PEGylation rate that we observed was due to changes in residue accessibility and not caused by an unforeseen effect the cations may have on the PEGylation reaction, we monitored the reaction rate of a position for which we observed no accessibility change in the structural analysis. A cysteine mutant at Ser436, positioned at the periphery of the transporter protein (Fig. 4a), exhibited minimal Na^+^-dependent accessibility changes (Fig. 4b, bottom row).

These accessibility measurements, along with our previous PEGylation results on three other VcINDY residues near the Na1 site (T154C, M157C and T177C, Supplementary Fig. 7e) ^29^, fully support the changes in protein dynamics predicted upon occupation of the Na1 and Na2 sites, and are consistent with our proposed induced-fit coupling model.

### Structural comparison of C_i_-Na^+^-S, Ci-Na^+^ and C_i_-*apo* states

The VcINDY structures determined in 300 mM Na^+^ and *apo* as reported here, together with previously-determined X-ray structure of the protein with both sodium and substrate bound ^1, 4^, allowed us to examine the structural changes of the transporter between the C_i_-Na^+^-S, C_i_-Na^+^ and C_i_-*apo* states. In addition to the flexibility observed in HP_in_b and TM10b, we observed amino acid sidechain movements both at the interface between the scaffold domain and the transport domain, as well as on the periplasmic surface of the protein.

At the scaffold – transport domain interface, side chains of several bulky amino acids rotated or shifted between the three states, including Phe100, His111 and Phe326 (Fig. 5a). On the periplasmic surface, Trp461 at the C-terminus is buried in the *apo* and Na^+^-bound structures (Fig. 5b). However, in the C_i_-Na^+^-S structure, the ring of the nearby Phe220 was rotated by ∼90°, pushing out the side chain of Trp461, leaving the C-terminus pointing to the periplasmic space. Accordingly, the loop between HP_out_b and TM10a moved into the periplasmic space of the apo VcINDY structure, displacing Glu394 and breaking its salt bridge with Lys337. Whereas no single switch was identified that can trigger conformational exchanges between the inward- and outward-facing states, local structural changes observed here suggest that small changes at multiple locations are required for inter-conformation transitions in VcINDY.

**Fig. 5.**
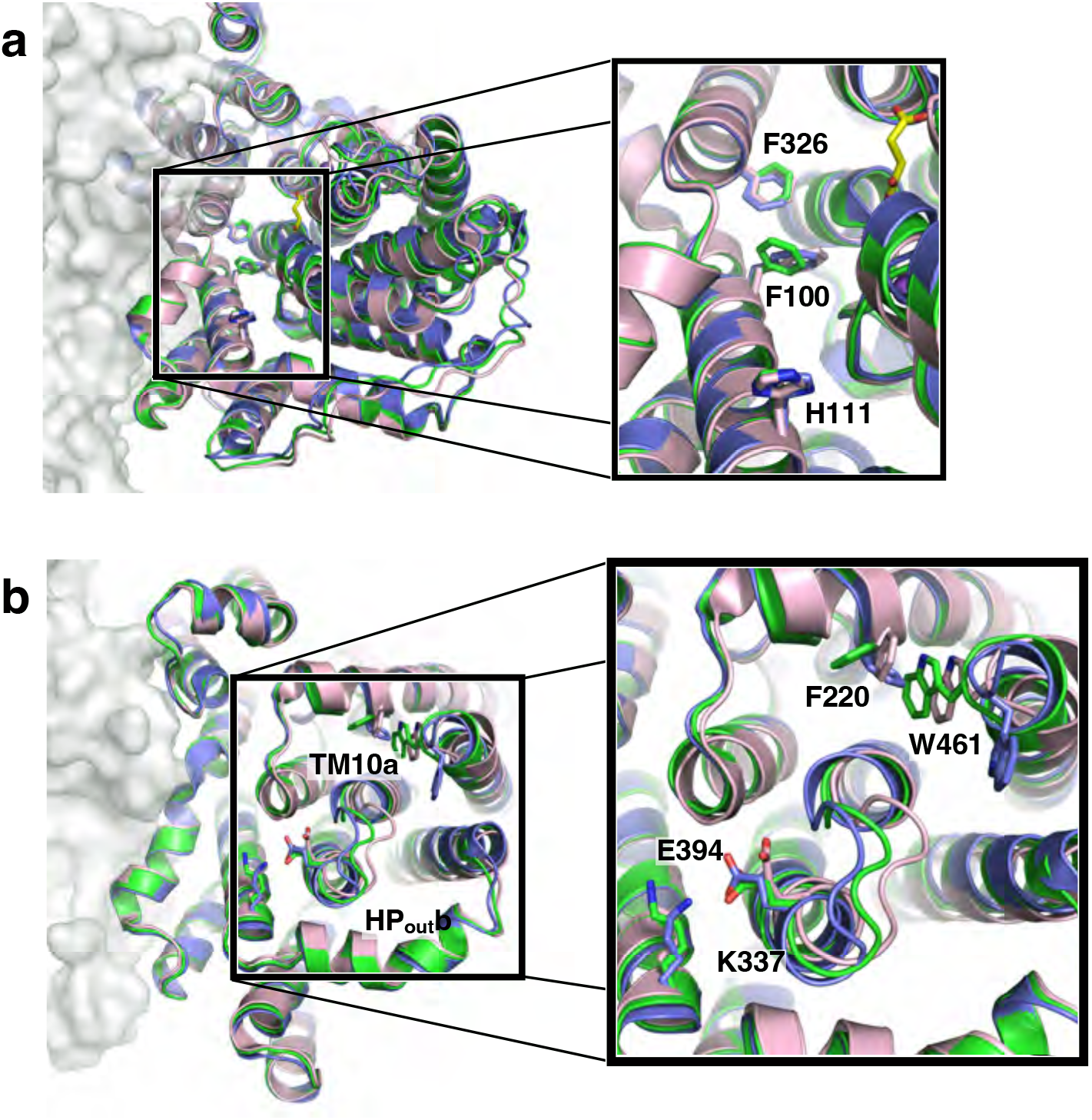
Movement of VcINDY’s amino acid side chains between its C_i_-Na^+^-S, C_i_-Na^+^ and C_i_-*apo* states. VcINDY structures in three states are overlaid: C_i_-Na^+^-S (blue), C_i_-Na^+^(green) and C_i_-*apo* (pink) states. **a**. At the scaffold-transport domain interface, the side chains of Phe100, His111 and Phe326 rotated between states. **b**. On the periplasmic surface, some loops and side chains move between the states, including Phe220, Lys337, Glu394 and Trp462.

In comparing maps of the three states, we noted the new VcINDY C_i_-Na^+^ map was sufficiently detailed to identify five ordered water molecules buried at the dimer interface (Supplementary Fig. 7c). The water molecules are not visible in previous maps, or the VcINDY-*apo* map, indicating the high-resolution of the new VcINDY C_i_-Na^+^ map was necessary for their identification. These waters are arranged in a square pyramidal configuration in the largely hydrophobic pocket, coordinated by only the symmetry related carbonyls of Phe92 and inter-water hydrogen bonds. The role of these waters in VcINDY folding or transport are unclear, though protein folding defects underlie several pathogenic mutations on the equivalent dimerization interface of NaCT ^5^.

## Discussion

Despite great advances in structural and mechanistic studies on membrane transporters over the past twenty years ^45–49^, the ion – substrate coupling mechanism is well characterized for only very few co-transporters, limiting our understanding of this fundamental aspect of the secondary-active transport mechanism. Here, we have described, for the first time, the structural basis of ion – substrate coupling for VcINDY, which reveals a novel induced-fit mechanism that ensures obligatory binding.

While Na^+^ sites in some other Na^+^-dependent transporters are buried in the middle of the protein ^37, 47, 48, 50^, the sodium sites in VcINDY are directly accessible from the extramembrane space. Previous experimental data support that Na^+^-driven DASS co-transporters operate via an ordered binding and release ^2, 25–29^. Specifically, Na^+^ binding occurs before substrate binding, while substrate release precedes Na^+^ release. For VcINDY, we have now observed that sodium release in the cytoplasm induces conformational changes between the C_i_-Na^+^ and C_i_-*apo* states, whereas the C_i_-Na^+^ and C_i_-Na^+^-S states are structurally similar (Fig. 6). Specifically, the movement of helices HP_in_b and TM10b is tightly coupled to Na^+^ binding. At the Na1 and Na2 sites, the sodium ions are stabilized via direct and ion — dipole interaction with the two helices. Therefore, upon Na^+^-release, the elimination of these interactions caused the relaxation of the HP_in_b and TM10b helices ^43, 44^. In the reverse reaction, the binding of Na^+^ ions, concurrent with helix re-ordering, creates a proper binding site for the substrate. Thus, we now have established a structural understanding of the Na^+^ — substrate coupling mechanism for this co-transporter. By extension, other DASS transporters may utilise a similar structural mechanism for Na^+^ — substrate coupling. (Fig. 6). Notably, this mechanism is distinct from the induced-fit proposed for the mitochondrial ADP/ATP exchanger, for which the protein changes are induced by the binding of the substrate itself ^51^.

**Fig. 6.**
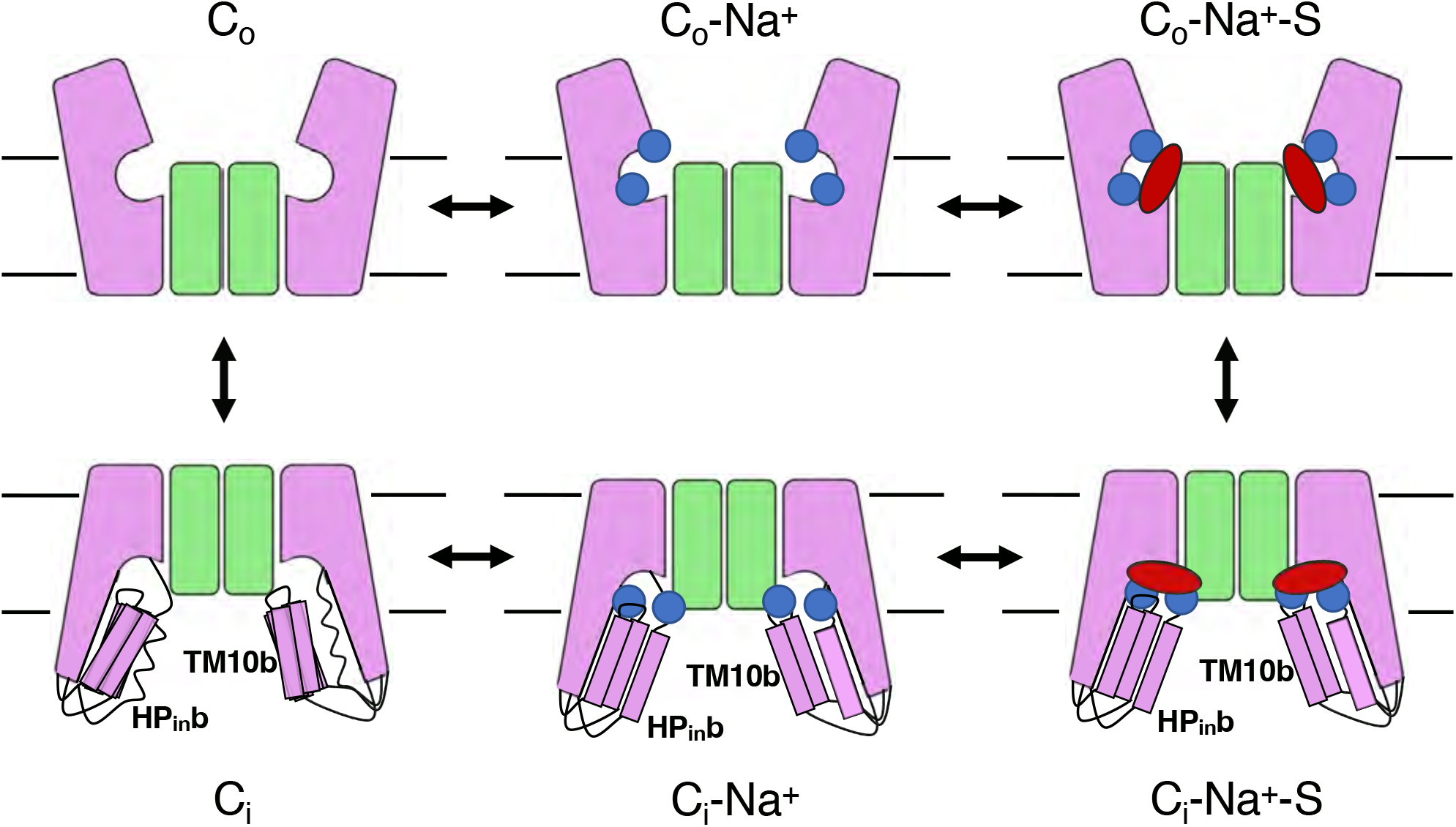
Schematic model of induced-fit mechanism for sodium — substrate coupling in VcINDY. In the absence of sodium ions, HP_in_b and TM10b are flexible. The binding of sodium ions (blue circles) organizes a proper binding site for the substrate, enabling the binding of succinate (red oval). The scaffold and transport domains in each protomer are colored as green and pink, respectively.

The structural basis of Na^+^-substrate coupling for VcINDY is also distinct from that of Glt_Ph_/Glt_Tk_, well characterized representatives of the dicarboxylate/amino acid:cation symporter (DAACS) family which otherwise share several commonalities with VcINDY: each transporter forms an oligomer with two re-entrant hairpin loops per protomer ^1, 4, 47^; each transporter co-transports its dicarboxylic substrate coupled to 3 Na^+^ ions ^22^, utilizing an elevator-like movement to pass the substrate across the membrane ^3, 52–56^ In addition, as we have shown here for VcINDY, an induced-fit mechanism has been suggested for both Glt_Ph_ and Glt_Tk_, which requires the initial binding of Na^+^ in order to prime the binding site for the substrate, aspartate ^37, 57^. However, the structural basis of Na^+^-substrate coupling in Glt_Ph_/Glt_Tk_ differs substantially from the coupling mechanism we observe for VcINDY. Rather than the general relaxation of a helix governing substrate binding site formation, the binding of Na^+^ to Glt_Ph_/Glt_Tk_ induces discrete conformational changes of a small number of amino acid residues centered on the highly conserved NMDGT motif. Among other conformational changes in this region, the Na^+^-induced rearrangement of sidechains in the NMDGT motif leads to the repositioning of a threonine sidechain, which is directly involved in aspartate binding, and an arginine, which moves to make space for the aspartate to bind, thus directly coupling Na^+^ binding to aspartate binding ^53, 57–59^. As is the case here for VcINDY (Fig. 6), the fully loaded and Na^+^-only bound structures of Glt_Ph_/Glt_Tk_ are largely identical ^52, 53, 57–59^, demonstrating that Na^+^ binding drives the formation of the substrate binding site, and not the substrate itself.

In addition to induced-fit, another mechanism for ion – substrate coupling of co-transporters has been proposed to be charge compensation ^5, 19^. Such a mechanism can greatly minimize the energy penalty for translocating charged substrates across the hydrophobic lipid bilayer ^60, 61^. Unlike for DASS exchangers ^19^, where charge compensation is the major force for overcoming the energy barrier in the C_o_ ←→ C_i_ transition, both local structural ordering and charge balance are needed for Na^+^-coupled co-transporters within the DASS family.

Comparison of the VcINDY structures reported here with those determined earlier ^1, 4, 19^, of three states in total, also sheds new light on the mechanism of the transporter’s conformational switching between the two sides of the membrane. As the C_i_-*apo* structure is significantly different from that of the C_i_-Na^+^-S state, their corresponding transitions to the outward-facing state: C_i_-*apo* to C_o_-*apo* and C_i_-Na^+^-S to C_o_-Na^+^-S, are different at the transport-scaffold domain interface (Fig. 6). Whereas the transition between C_i_-Na^+^-S and C_o_-Na^+^-S state can be described as rigid-body movement, as was seen in the DASS exchangers ^5^, the co-transporters’ C_o_-*apo* ←→ C_i_-*apo* state transition likely involves large structural rearrangements of the transport domain. Considering the pseudo-symmetry of the DASS fold, the C_i_-*apo* → C_o_-*apo* movement would require refolding of TM10b to pack against the scaffold domain, and possibly the concurrent unfolding of TM5b. This potential asymmetry between the *apo*-state transition (C_o_-*apo* ←→ C_i_-*apo*) and transition of the fully-loaded transporter (C_i_-Na^+^-S ←→ C_o_-Na^+^-S,) needs further investigation.

## Methods

### VcINDY expression and purification

Expression and purification of VcINDY was carried out according to our previous protocol ^1^. Briefly, *E. coli* BL21-AI cells (Invitrogen) were transformed with a modified pET vector ^62^ encoding N-terminal 10x His tagged VcINDY. Cells were grown at 32 °C until OD_595_ reached 0.8, protein expression occurred at 19 °C following IPTG induction, and cells were harvested 16 hrs post-induction. Cell membranes were solubilized in 1.2 % DDM and the protein was purified on a Ni^2+^-NTA column. For cryo-EM and substrate binding experiments, the protein was purified using size-exclusion chromatography (SEC) in different buffers. Protein used for the cysteine alkylation and transport assays was produced as described previously ^29^.

### Tryptophan fluorescence quenching assay

Tryptophan fluorescence quenching was used to measure affinity of succinate to purified VcINDY in detergent, using a protocol adapted from earlier work on other membrane transporters ^20, 30–33^. VcINDY purified by SEC in a buffer of 25 mM Tris Ph 7.5, 100 mM NaCl and 0.05% DDM was used to measure succinate affinity, while the 100 mM NaCl was replaced by 100 mM ChCl for affinity measurements in the absence of sodium. Protein was diluted to a final concentration of 4 μM in SEC buffer. Using a Horiba FluoroMax-4 fluorometer (Kyoto, Japan) at 22 °C and a 280 nm excitation wavelength, the emission spectrum was recorded between 290 and 400 nm. The emission maximum was determined to be 335 nm. Subsequently, the change in fluorescent emission at 335 nm was monitored with increasing concentrations of succinic acid (pH 7.5), from 0.1 μM to 1 mM. Each experimental condition was repeated 4 times. The binding curve was fit in Prism using a quadratic binding equation to account for bound substrate ^63^.

### Amphipol exchange and cryo-EM sample preparation

From Ni^2+^-NTA purified VcINDY, DDM detergent was exchanged to PMAL-C8 (Anatrace, Maumee, OH) as previously described ^19, 64^. Following further purification by SEC in buffer containing 25 mM Tris pH 7.5, 100 mM NaCl and 0.2 mM TCEP, the NaCl concentration was increased to 300 mM and the protein sample was concentrated to 1.3 mg/mL. For the *apo* protein preparation, NaCl in the abovementioned SEC buffer was replaced with 100 mM ChCl, and the protein sample was concentrated to 1.3 mg/mL.

Cryo-EM grids were prepared by applying 3 μL of protein to a glow-discharged QuantiAuFoil R1.2/1.3 300-mesh grid (Quantifoil, Germany) and blotted for 2.5 to 4 s under 100% humidity at 4 °C before plunging into liquid ethane using a Mark IV Vitrobot (FEI).

### Cryo-EM data collection

Cryo-EM data were acquired on a Titan Krios microscope with a K3 direct electron detector, using a GIF-Quantum energy filter with a 20-eV slit width. SerialEM was used for automated data collection ^65^. Each micrograph was dose-fractioned over 60 frames, with an accumulated dose of 65 e^-^/Å^2^.

### Cryo-EM image processing and model building

Motion correction, CTF estimation, particle picking, 2D classification, *ab initio* model generation, heterogenous and non-uniform refinement, and per particle CTF refinement were all performed with cryoSPARC ^66^. Each dataset was processed using the same protocol, except as noted.

Micrographs underwent patch motion correction and patch CTF estimation, and those with an overall resolution worse than 8 Å were excluded from subsequent steps. An ellipse-based particle picker identified particles used to generate initial 2D classes.

These classes were used for template-based particle picking. Template identified particles were extracted and subjected to 2D classification. A subset of well resolved 2D classes were used for the initial *ab initio* model building, while all picked particles were subsequently used for heterogeneous 3D refinement. After multiple rounds of 3D classification (*ab initio* model generation and heterogeneous 3D refinement with two or more classes), a single class was selected for non-uniform 3D refinement with C2 symmetry imposed, resulting in the final map.

All Cryo-EM maps were sharpened using Auto-sharpen Map in Phenix ^67^, models were built in Coot ^68^, and refined in Phenix real space refine ^69^. The model for VcINDY in NaCl was built using the structure of VcINDY embedded in a lipid nanodisc (PDB: 6WW5) as an initial model, with lipid and antibody fragments removed. The VcINDY model in choline used the structure of VcINDY in 300 mM NaCl, with ions and waters removed, as the starting model.

The NMR-style analysis used repeated runs of phenix.real_space_refine ^67^ to refine the models of VcINDY in *apo* and in 300 mM NaCl, with ions and waters removed, using unique computational seeds for each run. Each refinement was performed with simulated annealing, without NCS constraints or secondary structure restraints, and a refinement resolution limit of 3.23 Å for both maps. Analysis with or without map sharpening, or randomizing initial atomic positions using phenix.pdbtools, gave similar results.

### Cysteine alkylation assay

For the cysteine alkylation experiments, each purified cysteine mutant exchanged into reaction buffer containing 50 mM Tris, pH 7, 5% glycerol, 0.1% DDM and either 300 mM NaCl or 300 mM ChCl (Na^+^-free conditions). Protein samples were incubated with 6 mM mPEG5K and samples were taken at the indicated timepoints and immediately quenched by addition of SDS-PAGE samples buffer containing 100 mM methyl methanesulfonate (MMTS). Samples were analyzed with Coomassie Brilliant Blue-stained non-reducing polyacrylamide gels.

## Acknowledgements

This work was financially supported by the NIH (R01NS108151, R01GM121994 and R01-DK099023), the G. Harold & Leila Y. Mathers Foundation and the TESS Research Foundation (to D.N.W); and Wellcome Trust (210121/Z/18/Z) and BBSRC (BB/V007424/1) (to C.M.). D.B.S. was supported by the American Cancer Society Postdoctoral Fellowship (129844-PF-17-135-01-TBE) and Department of Defense Horizon Award (W81XWH-16-1-0153). We thank the following colleagues for helpful discussions: N. Coudray, R. Gonzalez Jr., M. Lopez Redondo, J.A. Mindell and E. Tajkhorshid. We are also grateful to the staff at the NYU Cryo-EM Facility and the NYU Microscopy Core for assistance in grid screening and the Pacific Northwest Center for Cryo-EM in data collection. EM data processing used computing resources at the HPC Facility of NYULMC.

## Author contributions

J.J.M. and J.S. and C.M. purified the proteins. J.J.M., J.C.S., and D.B.S. collected and analyzed the substrate binding data. C.M. did all the cysteine PEGylation experiments. D.B.S collected and processed the cryo-EM images and built the atomic models. D.B.S and D.N.W. analyzed the structures. D.B.S., C.M. and D.N.W. wrote the manuscript. All authors participated in the discussion and manuscript editing. C.M. and D.N.W. supervised the research.

## Data and Code Availability

Cryo-EM maps and models have been deposited in the Protein Data Bank and EMDB database, respectively, for VcINDY-Na^+^ (300 mM) (PBD ID: 7T9G; EMD-25757) and VcINDY-Ch^+^ (PBD ID: 7T9F; EMD-25756).

## Supplementary information

**Supplementary Fig. 1.**
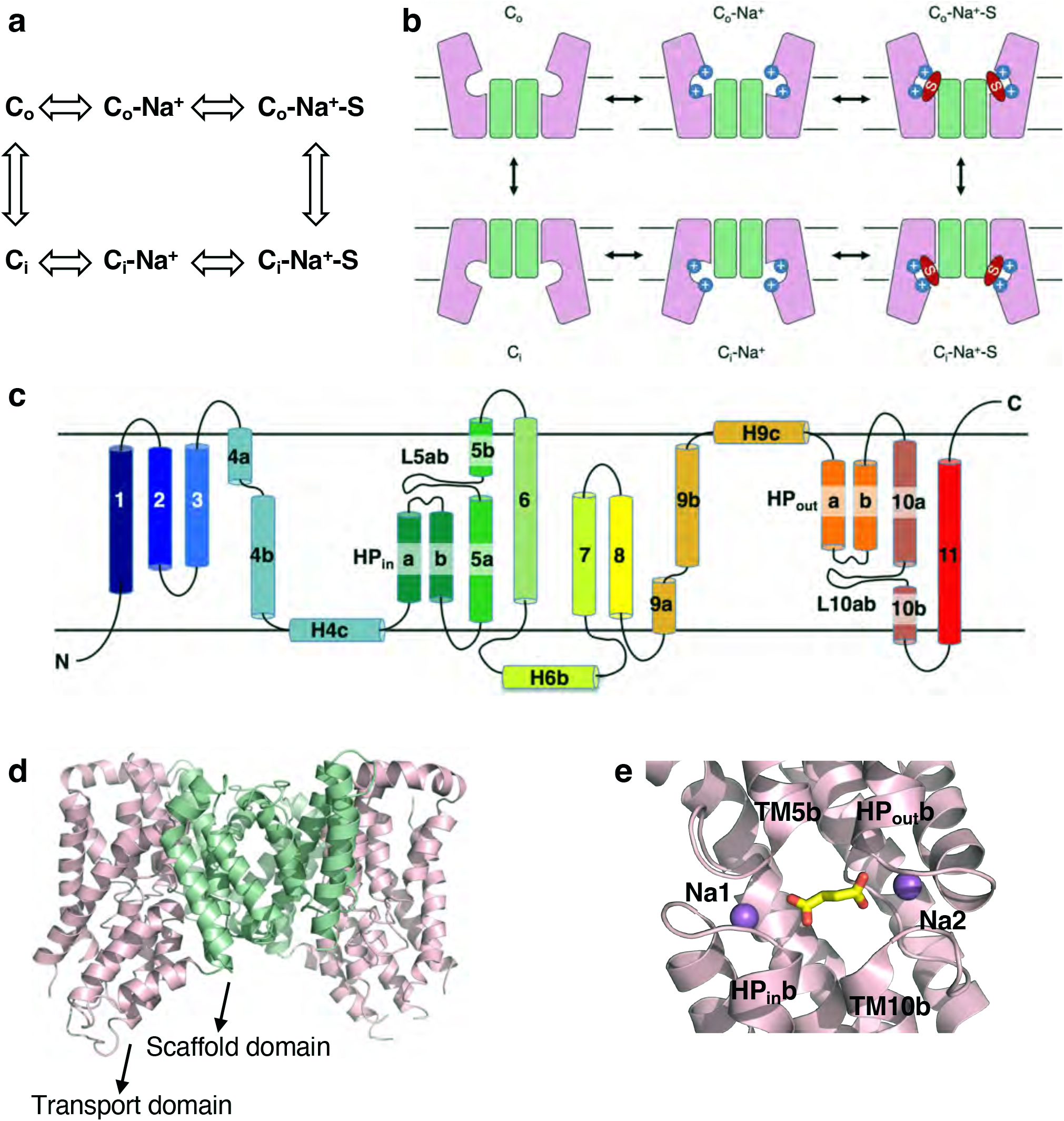
Kinetic cycle of VcINDY and molecular structure of its C_i_-Na^+^-S state. **a**, Kinetic cycle and, **b,** schematic model of VcINDY in its transport cycle. C_o_: outward-facing conformation; C_i_: inward-facing conformation; S: substrate. The number of co-transported Na^+^ for VcINDY is 3, but only two are shown here. All available biochemistry evidence indicates that sodium ions bind before and release after the substrate. **c,** Transmembrane topology of VcINDY. **d,** X-ray structure of VcINDY dimer. Each protomer consist of a scaffold domain and a transport domain. **e,** Structure of the substrate and sodium binding sites (PDB ID: 5UL7).

**Supplementary Fig. 2.**
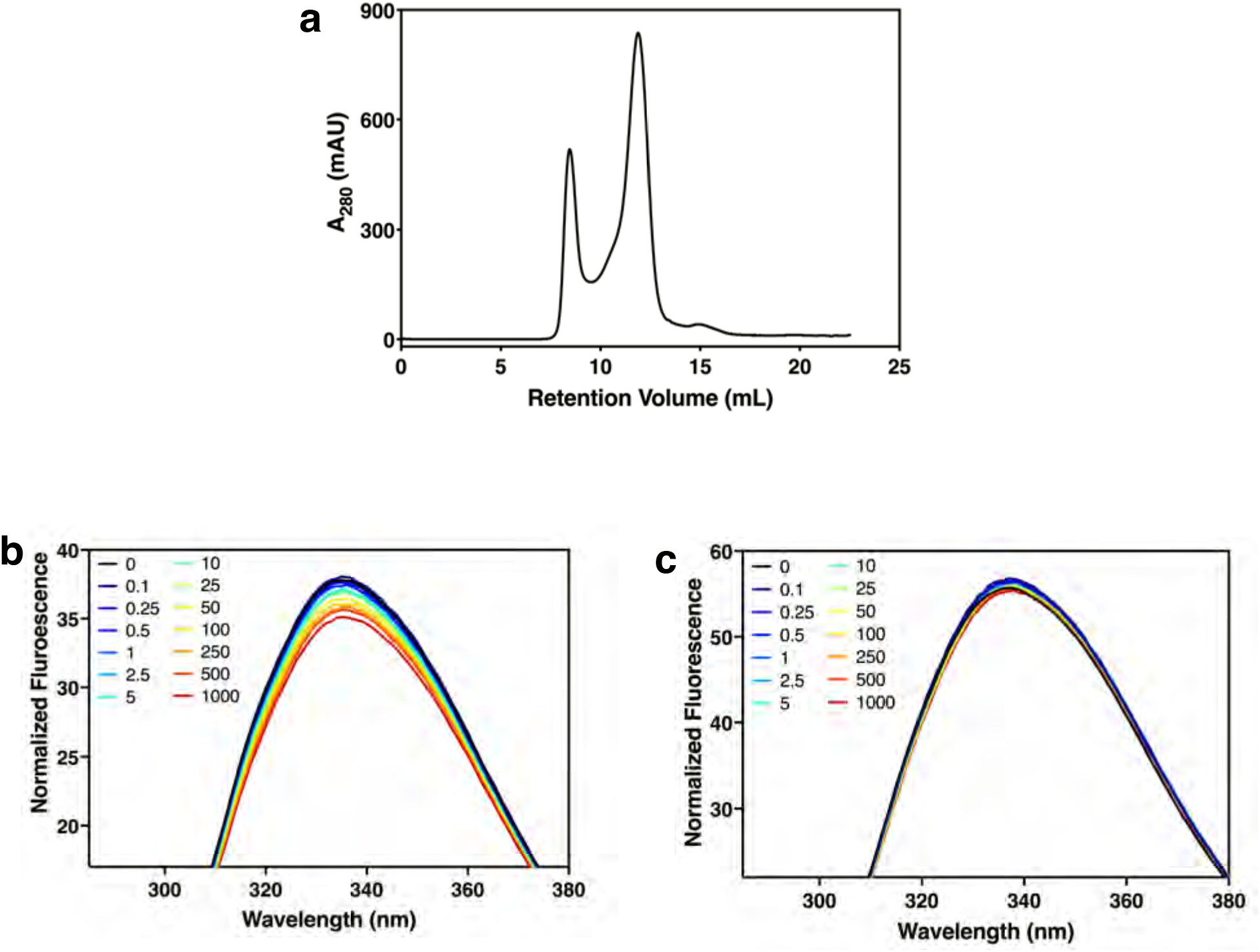
Purification and substrate binding measurements of VcINDY. **a,** Size-exclusion chromatography trace of VcINDY in dodecyl-maltoside detergent in the presence of 100 mM ChCl. Ch^+^ instead of Na^+^ was used as a cation to keep the protein stable. **b,** Measurements of succinate binding to detergent-purified VcINDY in the presence of 100 mM NaCl, using intrinsic tryptophan fluorescence quenching (*N* = 4). The *K*_d_ was determined to be 92.17± 47.38 μM. **c,** Measurements of succinate binding to VcINDY in the presence of 100 mM ChCl (*N* = 4). There was no binding that could be measured. In both **b** and **c**, the succinate concentrations in the legend are in μM.

**Supplementary Fig. 3.**
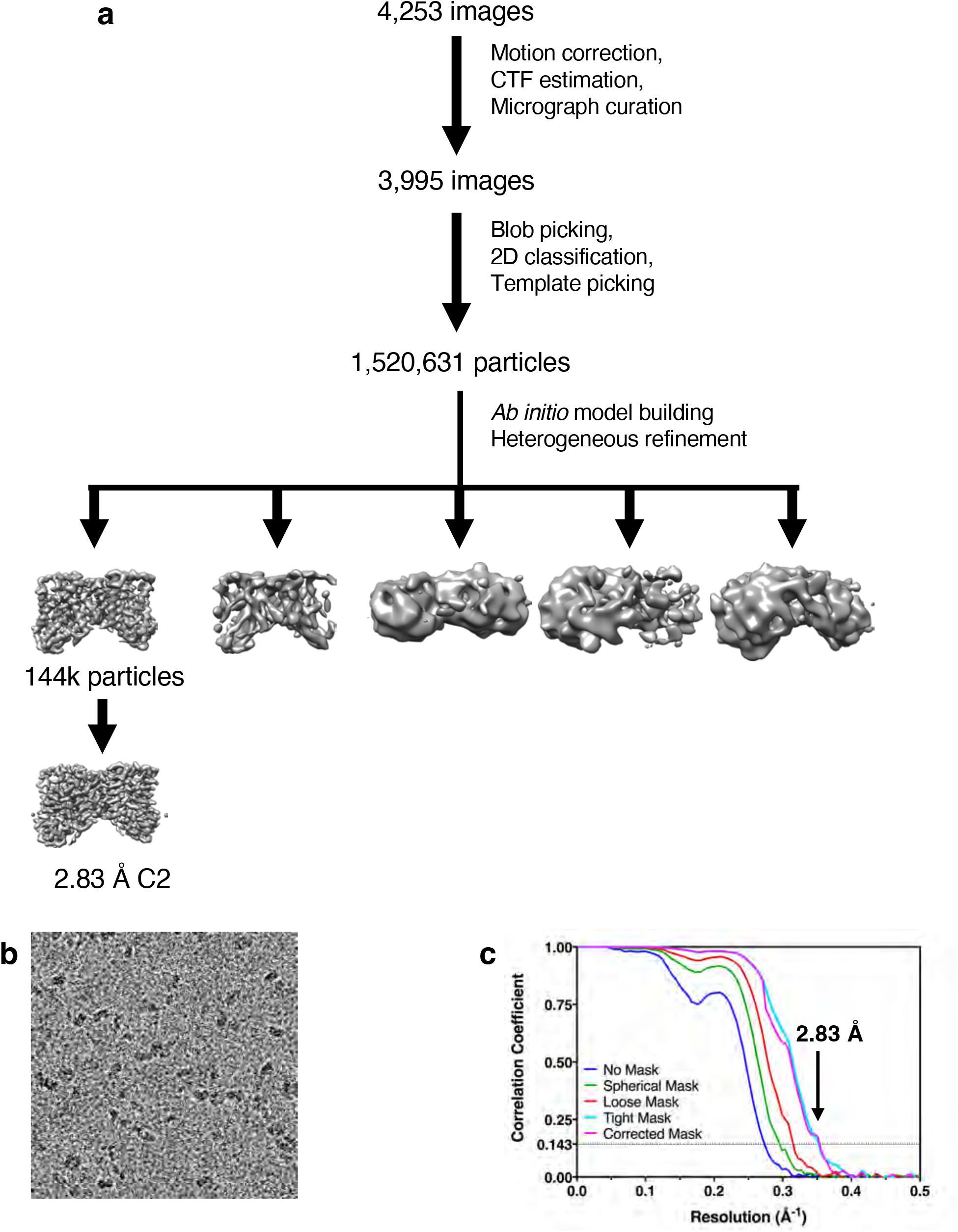
Cryo-EM structure determination of VcINDY in its C_i_-Na^+^ state solved in 300 mM Na^+^. **a,** Workflow of cryo-EM structure determination of VcINDY structure in 300 mM Na^+^. **b,** Cryo-EM micrograph. **c,** Fourier shell correlation curve. The gold-standard FSC resolution is indicated by the arrow.

**Supplementary Fig. 4.**
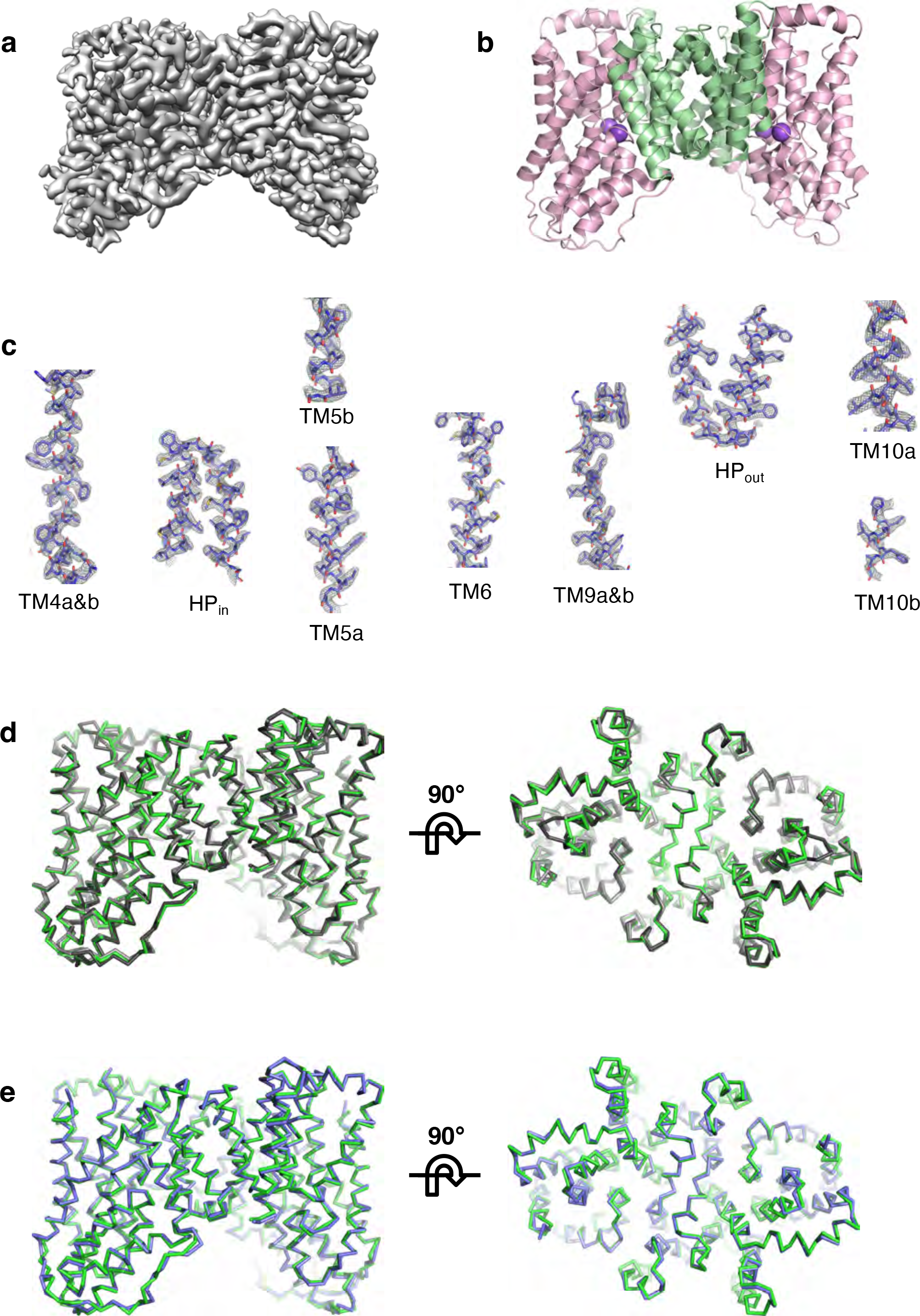
Cryo-EM structure of VcINDY in its C_i_-Na^+^ state solved in 300 mM Na^+^. **a,** Cryo-EM map at 2.83 Å resolution obtained in 300 mM NaCl. **b,** Model of VcINDY in its C_i_-Na^+^ state. The scaffold domain and the transport domain in each protomer are colored green and pink, respectively. **c,** Cryo-EM densities of individual helixes showing the quality of the model to map fitting. **d,** Overlay of the VcINDY structure determined in 300 mM Na^+^ (green) with two structures previously-determined in the presence of 100 mM Na^+^, with Fab bound in nanodisc (PDB ID: 6WW5; light grey) and without Fab but in amphipol (PDB ID: 6WU3; dark grey). Left, viewed from within the membrane plane. Right, viewed from the periplasmic space. **e,** Overlay of the VcINDY structure in the C_i_-Na^+^ state determined in 300 mM Na^+^ (green) with X-ray structure of VcINDY in the C_i_-Na^+^-S state (PDB ID: 5UL7; blue).

**Supplementary Fig. 5.**
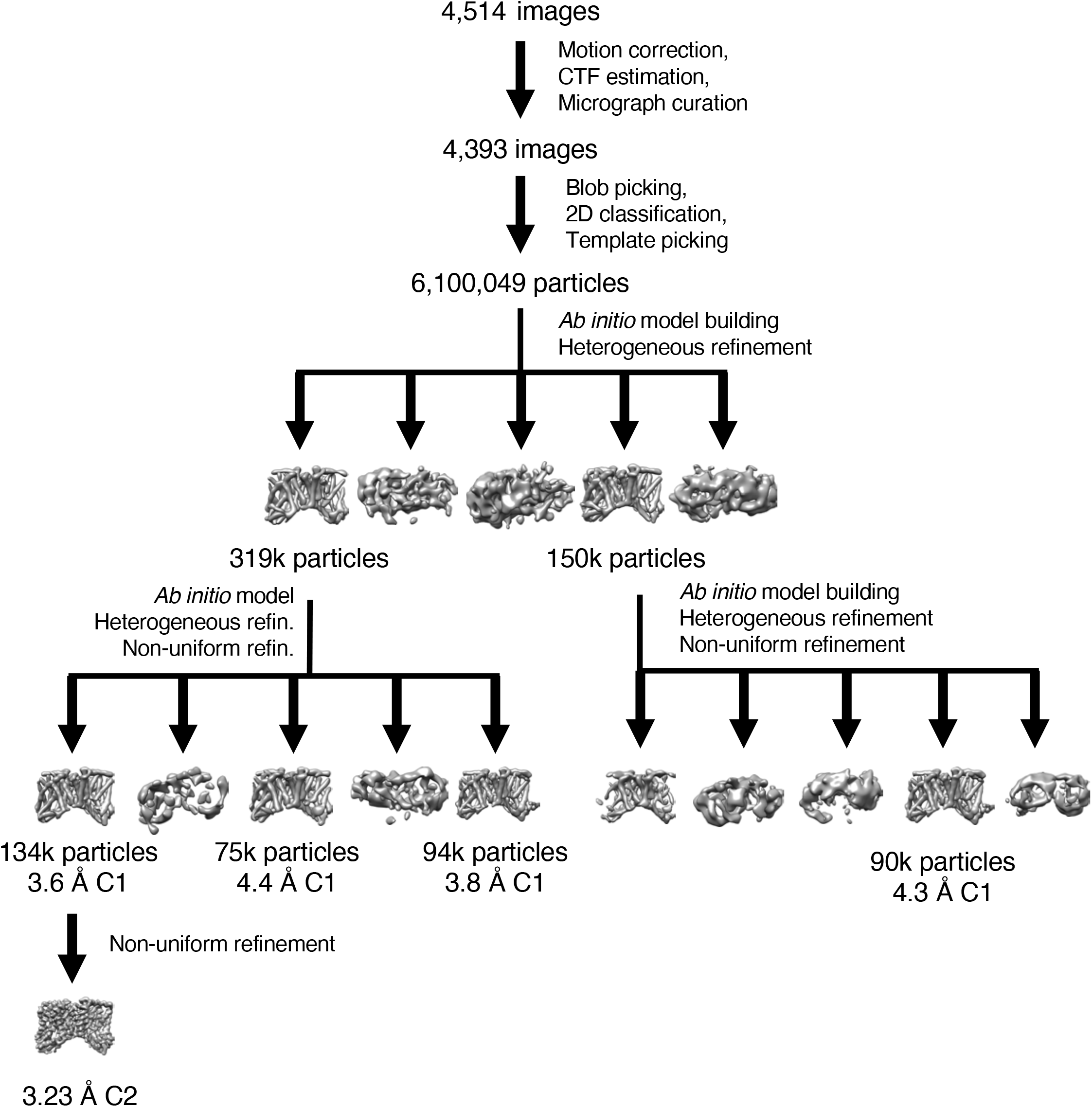
Workflow of cryo-EM structure determination of VcINDY in C_i_-*apo* state in choline.

**Supplementary Fig. 6.**
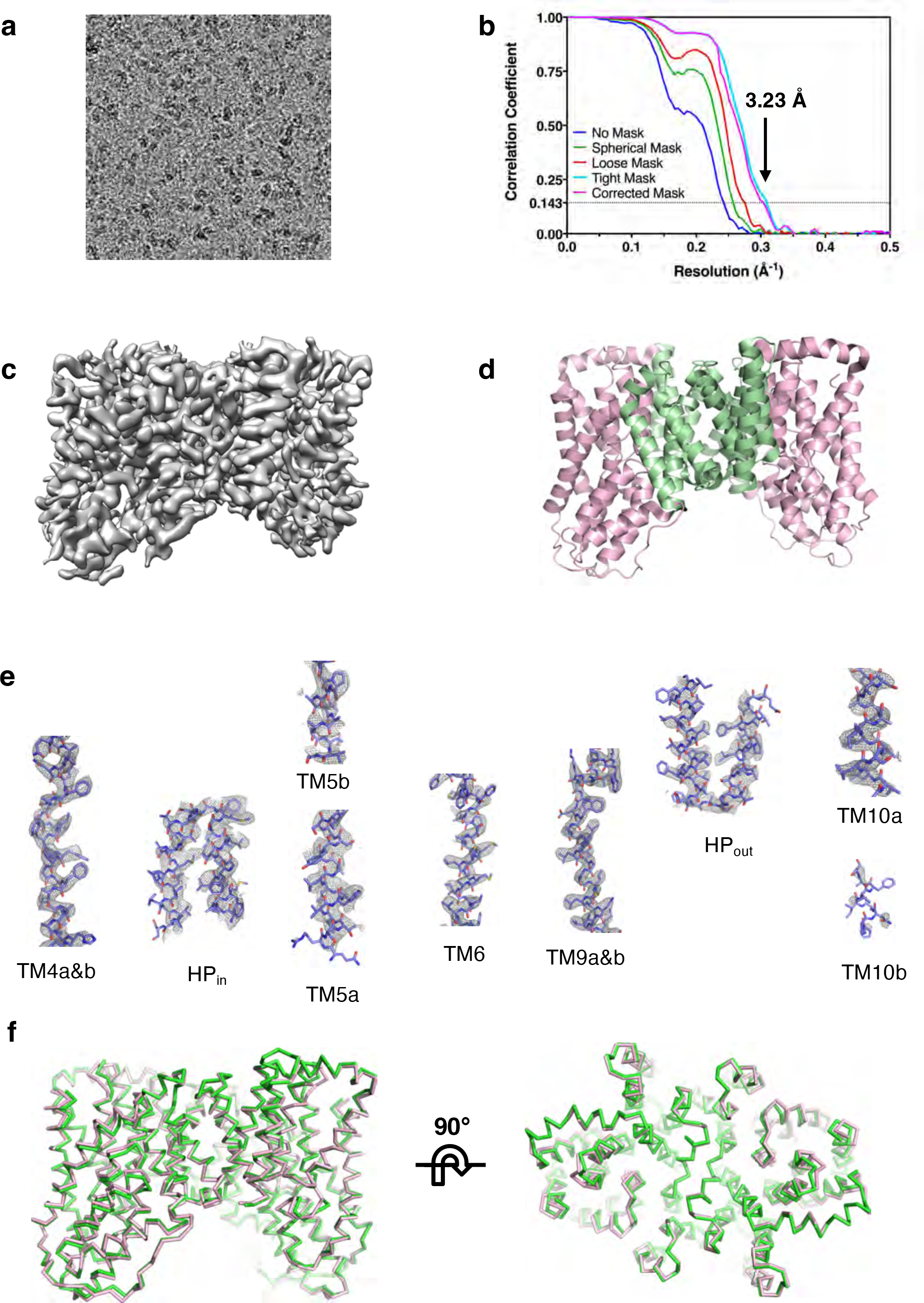
Cryo-EM structure determination of VcINDY in its C_i_-*apo* state solved in Ch^+^. **a,** Cryo-EM micrograph of VcINDY prepared in 100 mM Choline chloride. **b,** Fourier shell correlation curve. The gold-standard FSC resolution is indicated by the arrow. **c,** Cryo-EM map at 3.23 Å resolution obtained in Ch^+^. **d,** Model of VcINDY in its C_i_-*apo* state. The scaffold domain and the transport domain in each protomer are colored green and pink, respectively. **e,** Cryo-EM densities of individual helixes showing the quality of the model to map fitting. **f,** Overlay of the VcINDY structure in its C_i_-Na^+^ state determined in Na^+^ (green) with that of the C_i_-*apo* state determined in Ch^+^ (light pink). Left, viewed from within the membrane plane. Right, viewed from the periplasmic space.

**Supplementary Fig. 7.**
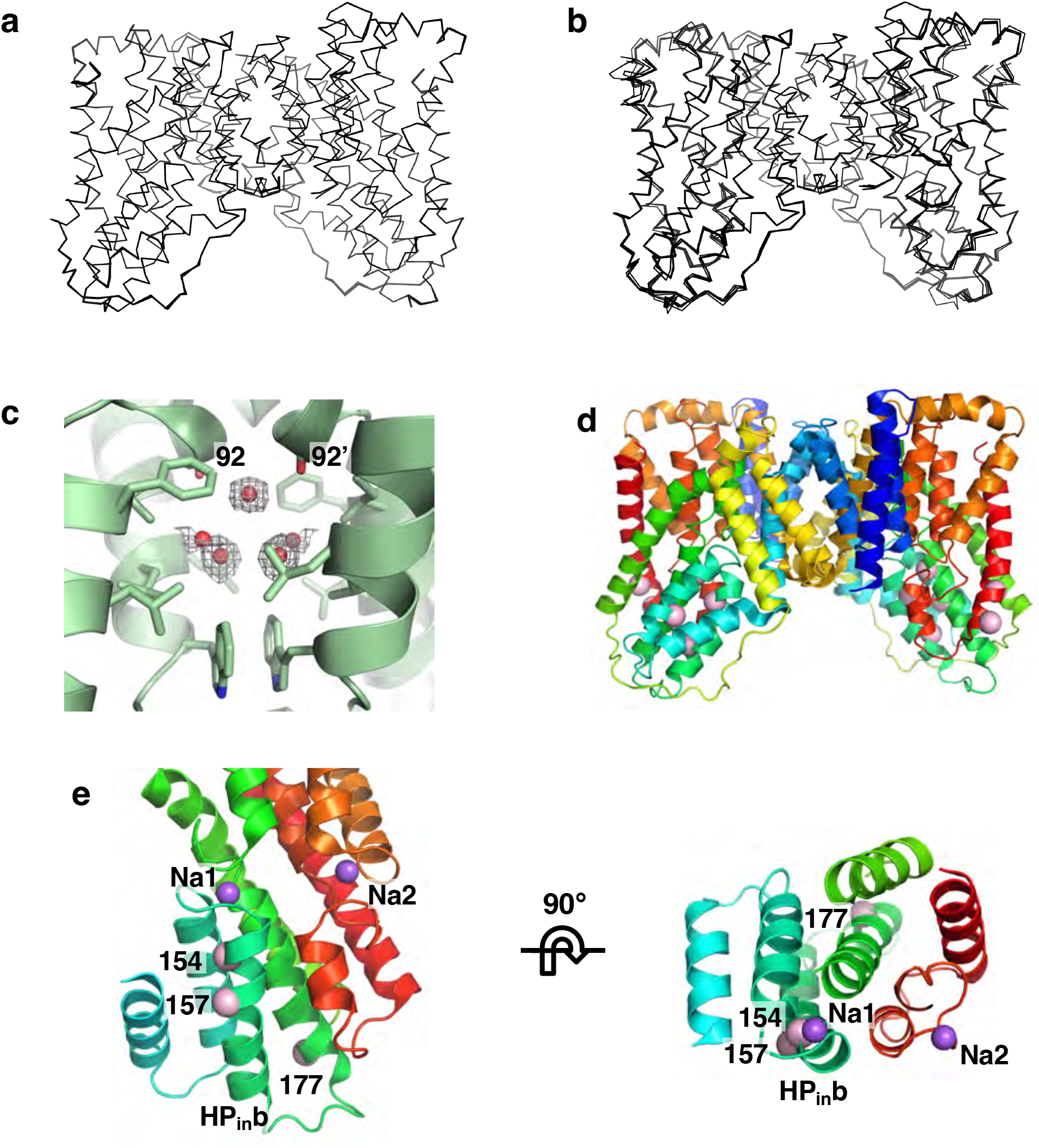
Flexibility of VcINDY-*apo* structure. **a,** NMR-style analysis of the VcINDY structure in Na^+^. **b,** Simulated annealing analysis of the VcINDY structure in choline^+^. The resolution limit for refinement in **a** and **b** was truncated to 3.23 Å. **c,** Water molecules observed in the VcINDY cryo-EM map in the C_i_-Na^+^ state. **d,** VcINDY dimer structure with positions of the residues labeled in the current work indicated by pink balls on the C_i_-Na^+^ state structure. **e,** Positions of relevant residues labeled in the previous work (Sampson, et al., J. Biol Chem, 2020, 295, 18524 – 18538).

**Supplementary Table 1.**
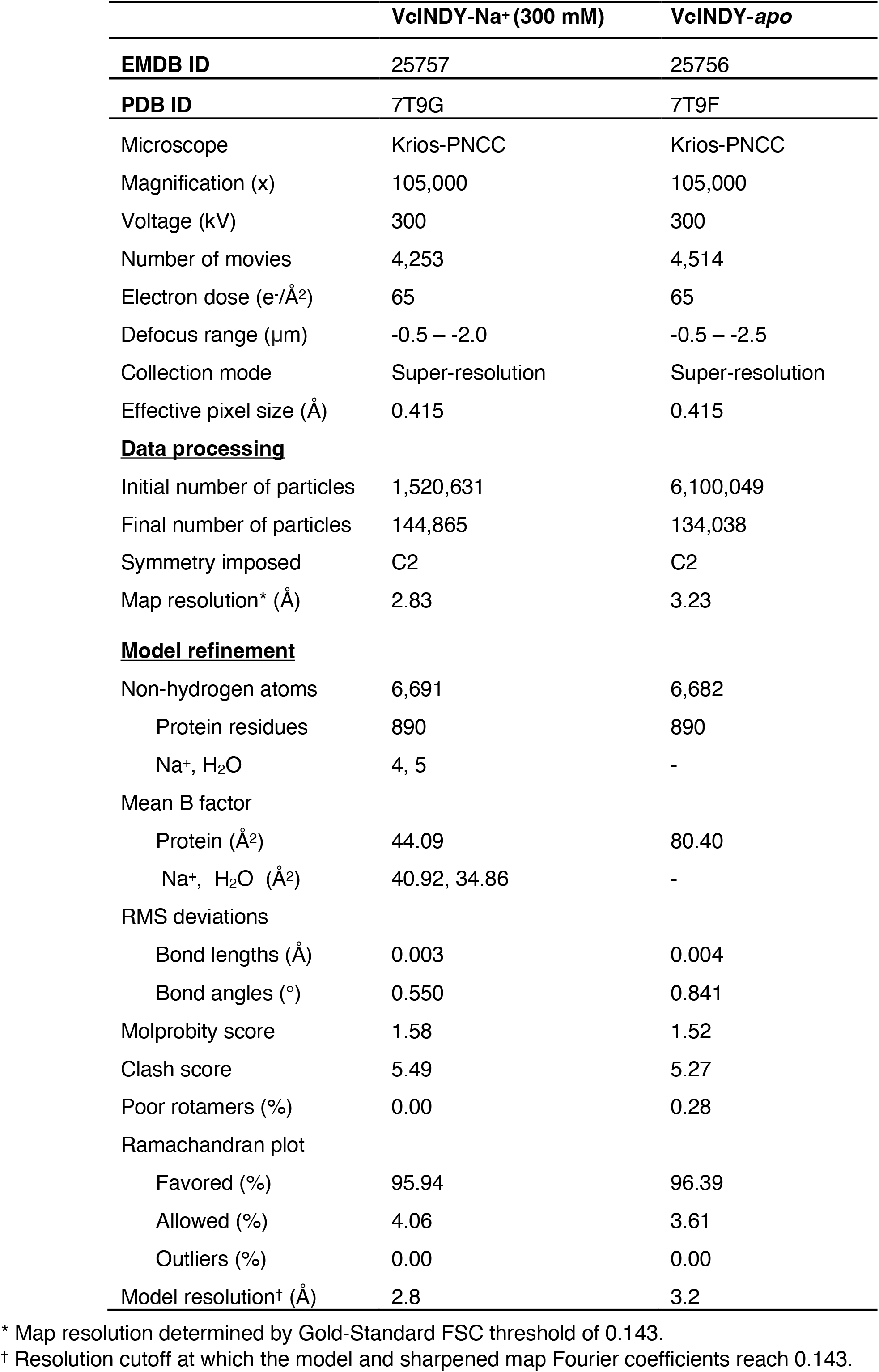
Cryo-EM data collection and structure determination of NaCT

